# Artificial selection footprints in domestic chicken genomes

**DOI:** 10.1101/2023.03.22.533830

**Authors:** Siwen Wu, Tengfei Dou, Kun Wang, Sisi Yuan, Shixiong Yan, Zhiqiang Xu, Yong Liu, Zonghui Jian, Jingying Zhao, Rouhan Zhao, Hao Wu, Dahai Gu, Lixian Liu, Qihua Li, Dong-Dong Wu, Zhengchang Su, Changrong Ge, Junjing Jia

## Abstract

Accurate and low-cost next generation sequencing technologies make re-sequencing of large populations of a species possible. Although many studies related to artificial selection signatures of commercial and indigenous chickens have been carried out, quite a small number of genes have been found to be under selection. In this study, we re-sequenced 85 individuals of five indigenous chicken breeds with distinct traits from Yunnan, a southwest province of China. By analyzing these indigenous chickens together with 116 individuals of commercial chickens (broilers and layers) and 35 individuals of red jungle fowl, we find a substantially large number of selective sweeps and affected genes for each chicken breed using a rigorous statistic model than previously reported. We confirm most of previously identified selective sweeps and affected genes. Meanwhile the vast majority (∼98.3%) of our identified selective sweeps overlap known chicken quantitative trait loci. Thus, our predictions are highly reliable. For each breed, we also identify candidate genes and selective sweeps that might be related to the unique traits of the chickens.

## Introduction

Chicken (*Gallus gallus*) has been domesticated by human for about 8,000 years (Lawal, et al. 2020), and multiple lines of evidence show that red jungle fowl (RJF) is the major ancestor of domestic chicken all over the world (Fumihito, et al. 1994; Lawal, et al. 2020; Wang, Thakur, et al. 2020). Artificial selection has resulted in numerous chicken breeds with distinct traits in different parts of the world for various purposes, including meat and egg production as well as recreation and ornament. Particularly, intensive systematic artificial selections carried out in a few companies in the last decays have led to highly production-efficient commercial broiler and layer lines used all over the world. Understanding the genetic basis of distinct traits of traditionally bred indigenous chicken as well as of commercialized broilers and layers is crucial to guide breeding programs to further improve the commercial lines and chicken welfare (Cheng 2010). Besides commercial lines, indigenous chicken breeds are also excellent model systems to study the relationships between genotypes and phenotypes (Burt 2007). Indeed, many studies have been done to reveal artificial selection signatures on commercial broilers and layers (Rubin, et al. 2010; Elferink, et al. 2012; Qanbari, et al. 2012; Fan, et al. 2013; Gheyas, et al. 2015; Boschiero, et al. 2018; Qanbari, et al. 2019) as well as on indigenous chickens (Guo, et al. 2016; Wang, et al. 2016; Wang, et al. 2017; Zhang, Yang, et al. 2017; Bortoluzzi, et al. 2018; Lawal, et al. 2018; Wang, Li, et al. 2020). These studies have identified genes or quantitative trait loci (QTLs) related to specific traits such as body size (Kerje, et al. 2003; McElroy, et al. 2006; Wang, et al. 2016; Wang, et al. 2017; Wang, Bu, et al. 2020; Wang, Cao, et al. 2020; Zhang, et al. 2020; Wang, Hu, et al. 2021), meat quality (Jennen, et al. 2005; Bihan-Duval, et al. 2018; Moreira, et al. 2018), egg production (Zhao, et al. 2021), feathering (Zhao, et al. 2016; Guo, et al. 2018; Zhang, et al. 2018; Yang, et al. 2019; Bortoluzzi, et al. 2020; Li, Lee, et al. 2020; Chen, et al. 2022; Qiu, et al. 2022), plumage color (Zhang, Liao, et al. 2017; Huang, Pu, et al. 2020; Huang, Otecko, et al. 2020; Nie, et al. 2021), skin color (Li, Sun, et al. 2020), behaviors (Luo, et al. 2020; Mehlhorn and Caspers 2020), immunity (Zou, et al. 2020), crest shapes (Li, Lee, et al. 2021), bone traits (Wang, et al. 2018; Li, Liu, et al. 2021; Kondoh, et al. 2022), rumpless trait (Noorai, et al. 2012; Freese, et al. 2014; Xu, et al. 2017; Wang, Khederzadeh, et al. 2021), and polydactyly (Zhang, et al. 2016; Chu, et al. 2017). However, genetic bases of many artificially selected traits, in particular, of indigenous chickens, are far from being fully understood. Yunnan, a southwest province of China, is one of the major centers where domestic chickens arise (Wang, Thakur, et al. 2020), and numerous chicken breeds have been raised in mountainous areas there. Among these indigenous chicken breeds are Daweishan, Hu, Piao, Wuding and Nine-claw chicken, each with distinct traits. Specifically, Daweishan chickens have a miniature body size (0.5∼0.8kg for female and 0.8∼1.2kg for male adults); Hu chickens have a large body size (3kg for female and 6kg for male adults) with extraordinarily stout legs; Piao chickens have a short tail (a rumpless phenotype); Wuding chickens have a relatively large body size with colorful feathers and thick fat; and Nine-claw chickens have nine claws with a middle-sized body.

To understand the domestication and genetic basis of the distinct traits of these indigenous chickens, we have re-sequenced 25 Daweishan chickens, 10 Hu chickens, 23 Piao chickens, 23 Wuding chickens and four Nine-claw chickens. By comparing the single nucleotide polymorphisms (SNPs) of these indigenous chicken populations with those of 35 RJF individuals as well as of 60 broiler individuals and 56 layer individuals, we were able to find more artificial selection signatures on the indigenous chickens, broilers and layers using a rigorous statistic model (Churchill and Doerge 1994) than previously reported (Rubin, et al. 2010; Gheyas, et al. 2015; Qanbari, et al. 2019). By comparing the selection signatures between the indigenous chicken breeds, RJF, broilers and layers, we have found numerous genomic regions and genes related to the breed-specific traits.

## Results

### Indigenous chicken breeds have higher nucleotide diversity

By using the re-sequencing short reads of 25 Daweishan chickens, 10 Hu chickens, 23 Piao chickens, 23 Wuding chickens, four Nine-claw chickens, 60 broilers, 56 layers and 35 RJFs, we detected 20, 16, 22, 19, 13, 17, 14 and 22 million single nucleotide variants (SNVs) and small indels in the eight chicken groups (Table 1), respectively, using the GRCg7b assembly as the template. After the redundant variants in the different groups were removed, there are 33 million variants totally. There are 17, 13, 19, 16, 11, 14, 12 and 19 million bi-allelic SNPs on autosomes and sex chromosomes in the groups of Daweishan, Hu, Piao, Wuding, Nine-claw chickens, broilers, layers and RJFs, respectively (Table 1). After removing the redundant ones in the different groups, we ended up with 26 million bi-allelic SNPs on autosomes and sex chromosomes, and our subsequent analyses will be focused on these bi-allelic SNPs in each chicken group (Table 1).

**Table 1:**
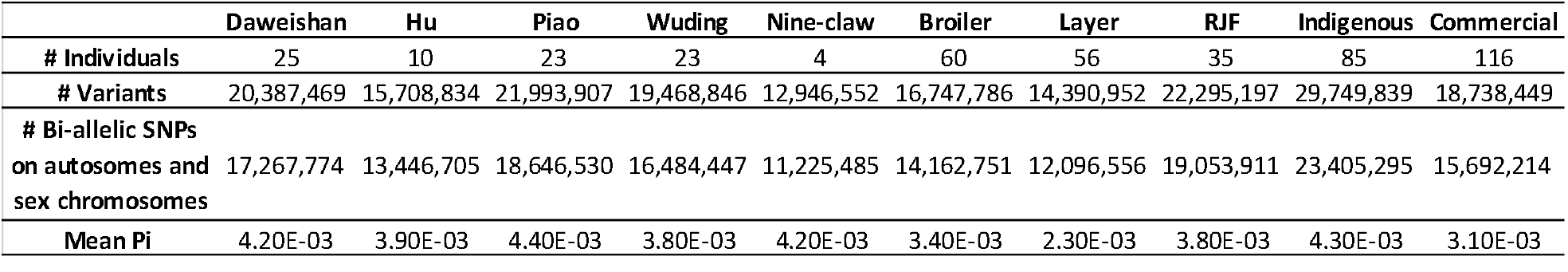
Summary of genetic variants in the chicken groups

When analyzing the mean nucleotide diversity (*π*) of each chicken group, we found the values of the broilers and the layers are smaller than those of the five indigenous chickens and RJFs (Table 1), indicating that the commercial lines are genetically more uniform than the indigenous lines and RJFs, as expected. The relatively low *π* values of commercial lines might be due to their close mating and linked selection (Qanbari, et al. 2019). At the same time, we detected 30 and 19 million SNPs on autosomes and sex chromosomes within the indigenous chicken group (85 individuals totally) and the commercial chicken group (116 individuals totally), respectively, of which, 23 and 16 million were bi-allelic SNPs for the two chicken groups, respectively (Table 1). Consistent with the aforementioned results, the mean *π* values of the indigenous group and the RJFs group are higher than that of the commercial group (Table 1), suggesting that artificial selection in commercial lines is more intensive than in indigenous chickens. Unexpectedly, the mean *π* value of the indigenous group is higher than that of the RJFs group with two different origins (India and Thailand, Materials and methods, Table 1), which might be due to some level of inbreeding on farms of the RJFs since their ancestors’ captures.

### Variants are enriched in non-coding regions while those in coding regions are largely tolerant

Based on the location of the bi-allelic SNPs, we classified them into seven categories, including intergenic (variants in intergenic regions), intronic (variants in introns), up/downstream (variants within a 1kb region up/downstream of transcription start/end sites), splicing (variants within 2 bp of a splicing junction), UTR 3’/UTR 5’ (variants in 3’/5’ untranslated regions), ncRNA (variants in non-coding RNA genes) and coding (variants in coding sequences). The relative abundances of the bi-allelic SNPs in each chicken group are shown in Table 2. Specifically, of the bi-allelic SNPs in each chicken group, 29.71%∼30.46% fall within intergenic regions, 50.44%∼51.57% are located in intronic regions, 2.86%∼2.97% fall within up/downstream regions, 4.08%∼4.24% are located in 3’ UTR/5’ UTR regions, 0.01% fall within splicing regions, 10.09∼10.18% are located in ncRNA regions, and 1.53%∼1.80% fall within coding regions (Table 2). Therefore, only a small portion (1.53%∼1.80%) of the bi-allelic SNPs fall within coding regions, while the vast majority (98.20%∼98.47%) are located in non-coding regions. As non-coding regions comprise 96.92% of the reference chicken genome (GRCg7b assembly), the SNPs are enriched in non-coding regions relative to in coding regions.

**Table 2:** Functional annotation of genetic variants in each chicken group

|  | Daweishan | Hu | Piao | Wuding | Nine-claw | Broiler | Layer | RJF | Indigenous | Commercial |
| --- | --- | --- | --- | --- | --- | --- | --- | --- | --- | --- |
| # SNPs in intergenic region (%) | 5,192,013 (30.07) | 3,995,687 (29.71) | 5,636,381 (30.23) | 4,963,186 (30.11) | 3,353,149 (29.87) | 4,237,766 (29.92) | 3,628,162 (29.99) | 5,803,793 (30.46) | 7,107,445 (30.37) | 4713,113 (30.03) |
| # SNPs in intronic region (%) | 8,802,652 (50.98) | 6,901,461 (51.32) | 9,465,663 (50.76) | 8,398,955 (50.95) | 5,788,439 (51.57) | 7,258,352 (51.25) | 6,201,733 (51.27) | 9,670,335 (50.75) | 11,805,669 (50.44) | 8,014,790 (51.07) |
| # SNPs in up/downstream (%) | 513,016 (2.97) | 398,721 (2.97) | 552,450 (2.96) | 489,473 (2.97) | 320,904 (2.86) | 419,495 (2.96) | 355,747 (2.94) | 548,631 (2.88) | 694,514 (2.97) | 464,953 (2.96) |
| # SNPs in UTR 3'/UTR 5' (%) | 719,457 (4.17) | 559,799 (4.16) | 778,556 (4.18) | 683,119 (4.14) | 458,168 (4.08) | 587,352 (4.15) | 495,415 (4.10) | 786,835 (4.13) | 992,584 (4.24) | 652,061 (4.16) |
| # SNPs in splicing region (%) | 1,166 (0.01) | 859 (0.01) | 1,258 (0.01) | 1,150 (0.01) | 662 (0.01) | 1,055 (0.01) | 866 (0.01) | 1,276 (0.01) | 1,791 (0.01) | 1,265 (0.01) |
| # SNPs in ncRNA (%) | 1,748,603 (10.13) | 1,368,611 (10.18) | 1,894,392 (10.16) | 1,672,022 (10.14) | 1,132,747 (10.09) | 1,428,434 (10.09) | 1,222,251 (10.10) | 1,935,153 (10.16) | 2,382,196 (10.18) | 1,587,018 (10.11) |
| # SNPs in coding region (%) | 290,867 (1.68) | 221,567 (1.65) | 317,830 (1.70) | 276,542 (1.68) | 171,416 (1.53) | 230,297 (1.63) | 192,382 (1.59) | 307,888 (1.62) | 421,096 (1.80) | 259,014 (1.65) |
| # Nonsynonymous-INTOL SNPs (%) | 14,069 (0.08) | 9,337 (0.07) | 16,330 (0.09) | 13,415 (0.08) | 6,452 (0.06) | 10,527 (0.07) | 8,594 (0.07) | 14,803 (0.08) | 57,713 (0.25) | 28,147 (0.18) |
| # Nonsynonymous-TOL SNPs (%) | 88,831 (0.51) | 66,095 (0.49) | 97,721 (0.52) | 84,157 (0.51) | 49,459 (0.44) | 69,943 (0.49) | 58,099 (0.48) | 93,118 (0.49) | 100,729 (0.43) | 64,301 (0.41) |
| # Synonymous SNPs (%) | 185,466 (1.07) | 144,341 (1.07) | 201,060 (1.08) | 176,605 (1.07) | 114,265 (1.02) | 147,808 (1.04) | 123,989 (1.02) | 197,449 (1.04) | 258,938 (1.11) | 164,231 (1.05) |
| # Stop-gain/loss (%) | 1,220 (0.01) | 862 (0.01) | 1,334 (0.01) | 1,145 (0.01) | 581 (0.01) | 962 (0.01) | 831 (0.01) | 1,290 (0.01) | 1,927 (0.01) | 1,169 (0.01) |
| # No prediction (%) | 1,281 (0.01) | 932 (0.01) | 1,385 (0.01) | 1,220 (0.01) | 659 (0.01) | 1,057 (0.01) | 869 (0.01) | 1,228 (0.01) | 1,789 (0.01) | 1,166 (0.01) |

Among the bi-allelic SNPs in coding regions of each chicken group, 33.33%∼38.33% (Daweishan = 35.71%, Hu = 34.55%, Piao = 36.47%, Wuding = 35.71%, Nine-claw = 33.33%, broiler = 34.97%, layer = 35.22%, RJF = 35.80%, indigenous = 38.33%, commercial = 36.36%) are amino acid-altering (AA-altering, i.e., nonsynonymous and stop-gain/loss) SNPs (Table 2). Among the nonsynonymous SNPs in each chicken group, most (Daweishan = 86.44%, Hu = 87.50%, Piao = 85.24%, Wuding = 86.44%, Nine-claw = 88.00%, broiler = 87.50%, layer = 87.27%, RJF = 85.96%, indigenous = 63.24%, commercial = 69.49%) are tolerant SNPs and only a small proportion (Daweishan = 13.56%, Hu = 12.50%, Piao = 14.76%, Wuding = 13.56%, Nine-claw = 12.00%, broiler = 12.50%, layer = 12.73%, RJF = 14.04%, indigenous = 36.76%, commercial = 30.51%) are intolerant, which might be harmful variants and thus under purifying selection in the chicken group (Table 2).

### Indigenous chicken breeds have a high portion of rare nonsynonymous SNPs

We compared the allele frequencies of the SNPs in coding regions in the groups of RJFs, indigenous and commercial populations. As shown in Figure 1a, all the three groups have higher portion of rare nonsynonymous SNPs than rare synonymous SNPs, indicating that rare nonsynonymous SNPs tend to be deleterious and thus under purifying selection. The same conclusion has been drawn in an earlier study for commercial chickens (Qanbari, et al. 2019). Interestingly, the indigenous chickens have the highest rare allele frequency densities for both nonsynonymous and synonymous SNPs, followed by RJFs and the commercial chickens. The earlier study also noted that RJFs had higher rare allele frequency densities than commercial chickens (Qanbari, et al. 2019). Intensive industrial selective breeding of commercial chickens can lead to the loss of rare alleles which might be slightly deleterious, thus the density of rare allele frequency of commercial chickens is the lowest among the three chicken groups. Consistent with our earlier results (Table 1), the indigenous chickens had higher rare allele frequency densities than RJFs. On the other hand, the indigenous chickens consist of five chicken breeds with distinct traits from different regions in Yunnan Province, China, thus they have higher nucleotide diversity (Table 1) compared with commercial chickens and RJFs, due to harboring more rare alleles (Figure 1a). Among the five different indigenous chicken breeds and two commercial chicken breeds, the density of the rare allele frequency of Piao chickens is the highest, and the value of layers is the lowest (Figure S1), consistent with their highest and lowest *π* values, respectively (Table 1).

**Figure 1.**
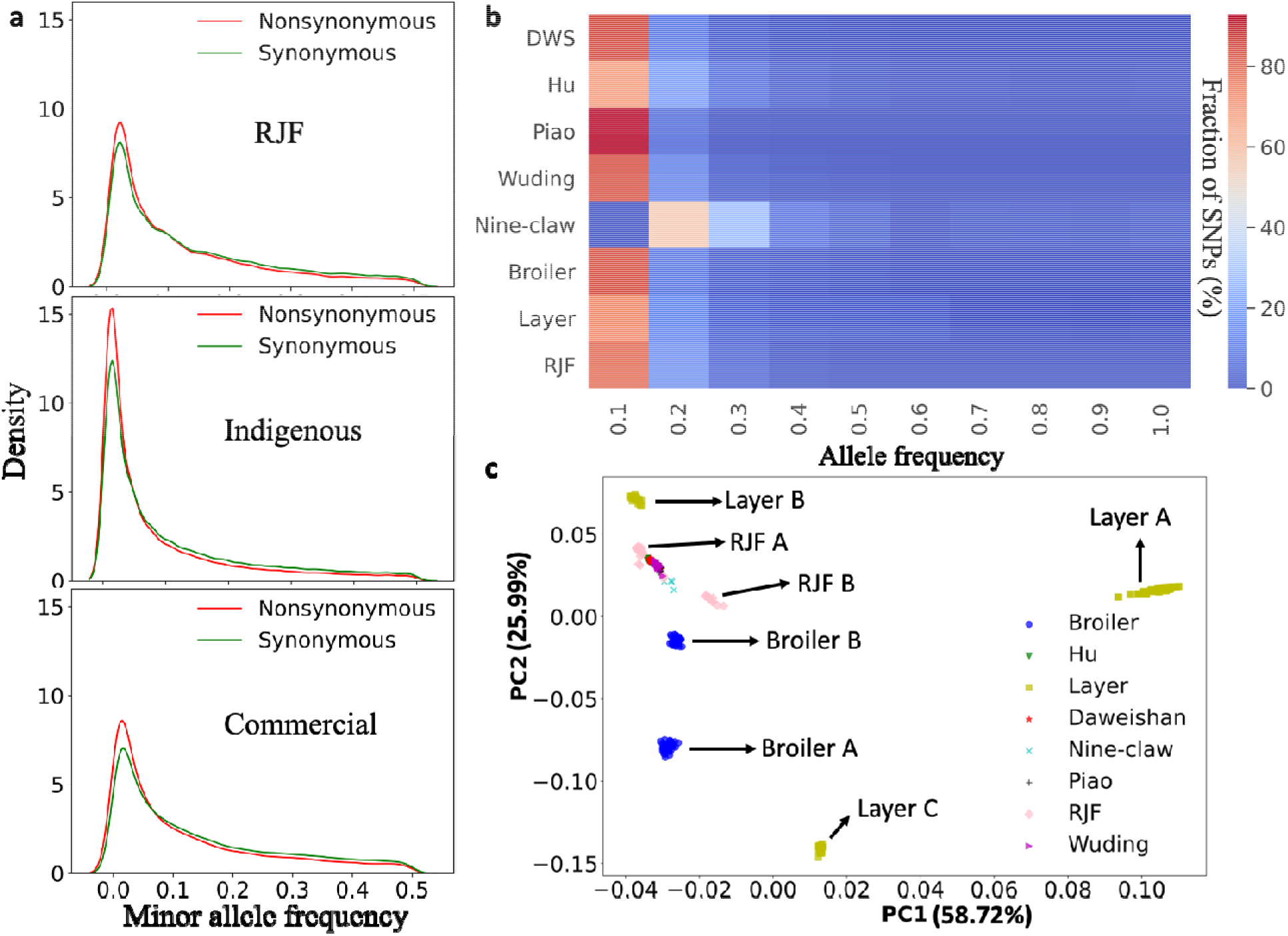
Analysis of frequency spectrums of SNPs. **a**. Distribution of minor allele frequency of synonymous and nonsynonymous SNPs in different chicken groups. **b**. Heatmap of allele frequency of group-specific SNPs. **c**. Principal component analysis of chicken population based on the detected 26 million SNPs.

### Only a small portion of breed-specific SNPs are fixed

We analyzed the group-specific SNPs in each chicken group, and found that RJFs have the highest number of unique SNPs (2.9 million) among the eight chicken groups (Table 3), which is consistent with the finding in the previous study (Qanbari, et al. 2019), suggesting a loss of ancestral alleles during chicken domestication. Except for Hu chicken and Nine-claw chicken with a small population size (Table 1), layers have the lowest number of unique SNPs (455k) and broilers have the second lowest number of unique SNPs (520k) among the eight chicken breeds (Table 3), while Daweishan, Piao and Wuding chickens have 1.1, 1.3 and 0.7 million unique SNPs, respectively, indicating once again that genetic diversity of indigenous chickens is higher than those of the layers and the broilers. From 0.83% (RJFs) to 1.39% (Wuding chicken) of the group-specific SNPs are missense mutations (Table 3). Most of the group-specific SNPs have allele frequencies lower than 0.5, and only a very small portion (0.05%∼0.59%) are close to being fixed (allele frequency > 0.9) in all the eight groups of chickens (Table 3). The same is true for the missense SNPs (0.04%∼0.48%) (Table 3). More details of the group-specific missense SNPs and affected genes are listed in Tables S1∼S8.

**Table 3:**
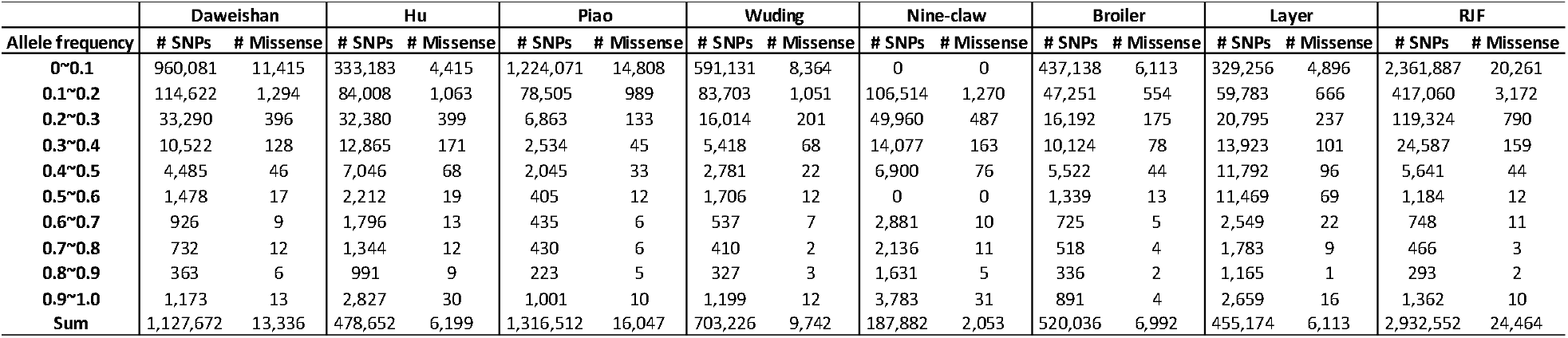
Frequencies of group-specific SNPs in the eight groups of chickens

We also compared allele frequency spectrums of the group-specific SNPs in the eight chicken groups (Figure 1b). Group-specific alleles of the layers tend to have higher frequencies than those in other groups (except for Hu chicken and Nine-claw chicken with a small population size), consistent with the finding in the earlier study (Qanbari, et al. 2019). This might be a result of artificial selection which increase the frequency of favorable alleles in layers.

### The indigenous chickens are more closely related to one another while commercial chickens are genetically different

To reveal the genetic relationships of the individual chickens, we performed a principal component analysis (PCA) based on occurring patterns of the ∼26 million bi-allelic SNPs. As shown in Figure 1c, the five indigenous breeds from Yunnan Province, China, are clustered together with RJFs from northern Thailand (RJF A) that is geographically close to Yunnan Province, China, while RJFs from India (RJF B) form a separate cluster nearby. This result suggests that the five indigenous breeds are closed related to one another, and they are also more closely related to RJFs from northern Thailand (RJF A) than to those from India (RJF B). On the other hand, brown egg layers (Layer B) and broilers from Indian River International (Broiler B) form two tight clusters close to the cluster of indigenous chickens and RJF A, while white egg layers (Layer A), crossbred layers (Layer C) and broilers from France (Broiler A) form clusters far away from the cluster of indigenous chickens and RJF A. These results suggest that broilers and layers with different origins have quite different genetic structures although they might have similar productivities.

### A rigorous Null model facilitates sensitive detection of selective sweeps

To detect selection signatures of each chicken breed, we identified selective sweeps (Sabeti, et al. 2006) along the chromosomes based on the frequencies of the bi-allelic SNPs. We used both genetic differentiation (*F*_*ST*_) and nucleotide diversity (*π*) to determine the selective sweeps of each chicken breeds. A sliding window of 40 kb with 20 kb step size was used to compute both *F*_*ST*_ and *π*. To identify selective sweeps more sensitively, unlike previous studies (Qanbari, et al. 2019) that used the sample mean and standard deviation to compute Z-values, we computed *ZF*_*ST*_ and *Z*|*Δπ*| values for each comparison based on the mean and standard deviation of Null models generated by permutating allele frequencies of the samples (Churchill and Doerge 1994) (Materials and methods). We consider a window with *ZF*_*ST*_ > 6 or *Z*|*Δπ*| > 3.09 (P-value < 0.001) as a selective sweep. Since adjacent windows can overlap with each other, we merged the overlapping selective sweep windows to form a discrete selective sweep (DSS) (Materials and methods). To find selection signatures of the chicken groups, we conducted a total of 19 different comparisons (Tables 4 and S9). The selective sweep windows identified by either of the two methods are distributed along all the chromosomes with varying densities (Figures 2, 3, S2 and S3). We generally detected more DSSs using *ZF*_*ST*_ (806∼2,125 DSSs) than using *Z*|*Δπ*| (110∼818 DSSs) for all the 19 comparisons (Tables 4 and S9) even using a higher *ZF*_*ST*_ cutoff, suggesting that *ZF*_*ST*_ is more sensitive than *Z*|*Δπ*| to identify selective sweeps. However, less than 60% (9.16%∼58.23%) of the DSSs identified by *Z*|*Δπ*| overlap with those identified by *ZF*_*ST*_ (Tables S10∼S28) in the 19 comparisons, indicating that the results of the two methods are largely complementary with each other. We thus take their union as the final prediction of DSSs (Tables S10∼S28). We finally identified 1,073∼2,745 DSSs consisting of 1,998∼7,284 windows containing 528∼2,147 genes in each of the 19 comparisons (Tables 4 and S9). Therefore, we find much more selective sweeps than the previous study using a mixture model (∼60 selective sweep windows of 40 kb size) (Qanbari, et al. 2019). The DSSs in the 19 comparisons have a varying length ranging from 40 kbp to 2,240 kbp with a median length of 60 kbp, and 91.71% of them are shorter than 140 kbp (Figure 4a). The total length of the DSSs in each comparison consist of 5.85% (Nine Claw VS RJF) ∼ 18.88% (Broiler VS Layer) of the reference genome (GRCg7b assembly) (Figure 4b). In general, comparisons with broilers alone as one of the two compared groups tend to have a high portion of genome under selection (Figure 4b), suggesting that broilers have gone through most extensive selection.

**Table 4:** Summary of putative selective sweeps and DSSs found in each chicken group in comparison with the RJFs

| Breeds | # Analyzed windows |  | # Windows passed critical cut-off |  | # Discrete selective sweeps (# genes) |  | # Final putative selective sweep windows (# DSSs, # genes in DSSs) |
| --- | --- | --- | --- | --- | --- | --- | --- |
|  | ZF<sub>ST</sub> | Z\|ΔPi\| | ZF<sub>ST</sub> > 6 | Z\|ΔPi\| > 3.09 | ZF<sub>ST</sub> > 6 | Z\|ΔPi\| > 3.09 |  |
| Daweishan VS. RJF | 51,857 | 51,789 | 4,157 | 595 | 1,268 (978) | 263 (790) | 4,619 (1,550, 1706) |
| Hu VS. RJF | 51,822 | 51,719 | 3,407 | 1,601 | 1,054 (783) | 645 (802) | 4,415 (1,811, 1,425) |
| Piao VS. RJF | 51,860 | 51,789 | 4,057 | 383 | 1,313 (845) | 152 (647) | 4,381 (1,483, 1,455) |
| Wuding VS. RJF | 51,853 | 51,782 | 3,752 | 1,554 | 1,255 (811) | 764 (892) | 4,962 (2,077, 1,617) |
| Nine-claw VS. RJF | 51,769 | 51,615 | 1,640 | 497 | 806 (398) | 262 (176) | 1,998 (1,073, 528) |
| Broiler VS. RJF | 51,643 | 51,756 | 6,366 | 2,150 | 1,784 (1,477) | 818 (646) | 7,012 (2,745, 1,661) |
| Layer VS. RJF | 51,808 | 51,725 | 3,808 | 791 | 1,521 (896) | 407 (404) | 4,192 (1,926, 1,139) |
| Indigenous VS. RJF | 51,886 | 51,793 | 4,294 | 301 | 1,183 (817) | 130 (528) | 4,572 (1,319, 1,333) |
| Commercial VS. RJF | 51,658 | 51,767 | 5,063 | 1,146 | 1,787 (1,191) | 544 (584) | 5,540 (2,382, 1,544) |

**Figure 2.**
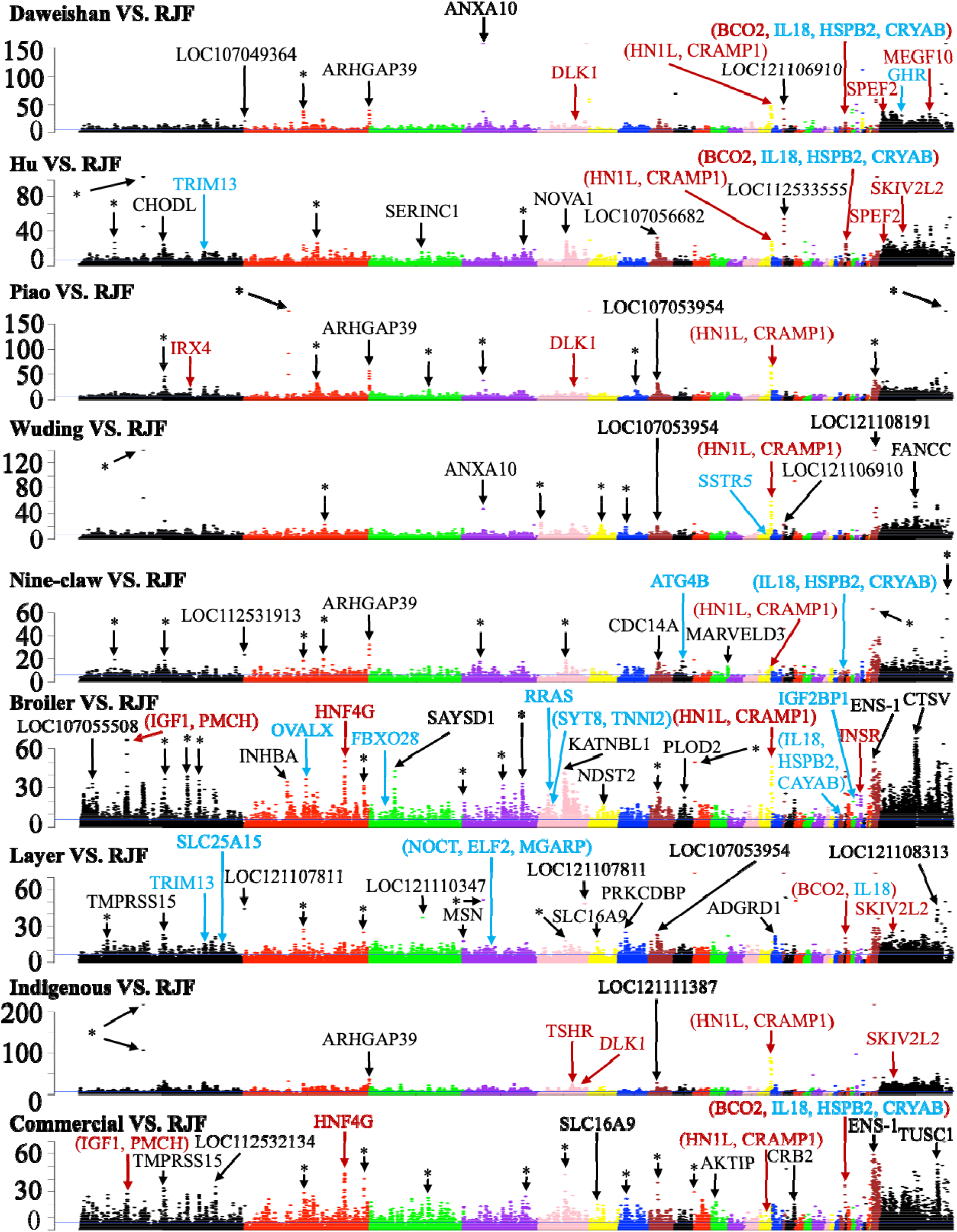
Manhattan plots of values of each window on each chromosome for the indicated comparisons. The blue horizontal line indicates the cutoff = 6. Examples of genes in significant selective sweep windows are shown in different color. Genes that have been previously reported in selective sweep windows are shown in red, genes in our predicted selective sweep windows potentially related to the specific traits of each chicken breed are shown in blue, and genes in novel selective sweep windows with extremely high *ZF*_*ST*_ values are shown in black. Asterisk represents selective sweep windows lacking annotated genes.

**Figure 3.**
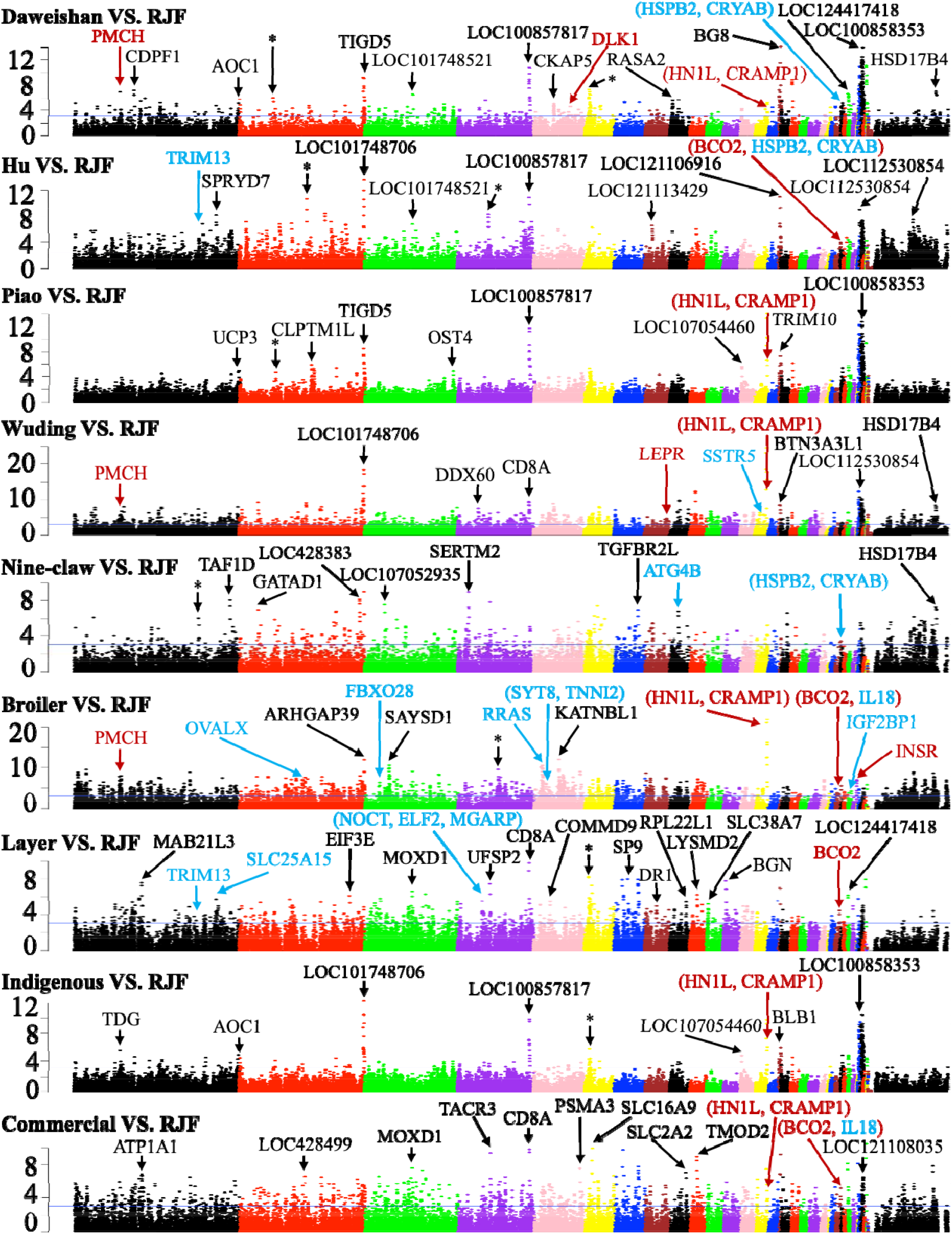
Manhattan plots of values of each window on each chromosome for the indicated comparisons. The blue horizontal line indicates the cutoff = 3.09. Examples of genes in significant selective sweep windows are shown in different color. Genes that have been previously reported in selective sweep windows are shown in red, genes in our predicted selective sweep windows potentially related to the specific traits of each chicken breed are shown in blue, and genes in novel selective sweep windows with extremely high *Z*|*Δπ*| values are shown in black. Asterisk represents selective sweep windows lacking annotated genes.

**Figure 4.**
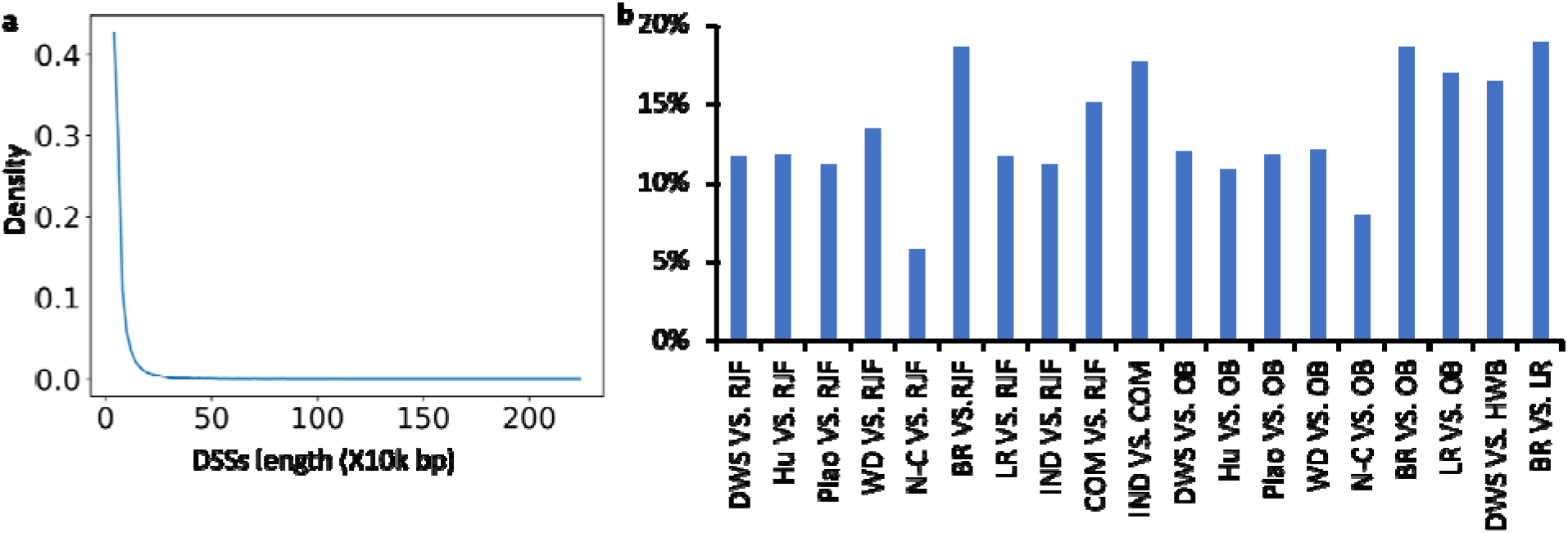
Summary of the DSSs lengths. **a**. Distribution of the lengths of the DSSs pooled from the 19 comparisons. **b**. Ratio of the DSSs lengths in each comparison with respect to the length of the reference genome (GRCg7b assembly). Abbreviations: DWS for Daweishan, WD for Wuding, N-C for Nine-claw, BR for Broiler, LR for Layer, IND for indigenous, COM for commercial, OB for Other Breeds and HWB for Hu+Wuding+Broiler.

### The 19 comparisons reveal selection signatures of the chicken groups

Firstly, to find the genetic differences between artificial selection and natural selection, we compared each chicken breed (including indigenous chicken group and commercial chicken group) with the RJFs. As summarized in Table 4 and Tables S10∼S18, we identified varying numbers of selective sweeps (1,998∼7,012) and DSSs (1,073∼2,745) involving 528∼1,706 genes for the eight comparisons, suggesting that these chicken breeds might have gone through different levels of artificial selection. For example, the Broiler VS. RJF comparison yields the highest numbers of selective sweeps (7,012) and DSSs (2,745), suggesting again that broilers might have gone through the most intensive artificial selection. Among the five indigenous breeds, Wuding chickens might have gone through the most intensive artificial selection with the highest numbers of selective sweeps (4,962) and DSSs (2,077). In addition, we identified more selective sweeps (5,540) and DSSs (2,382) for the Commercial VS. RJF comparison than those (4,572 selective sweeps and 1,319 DSSs) in Indigenous VS. RJF comparison (Table 4), suggesting that the commercial chickens have gone through more intensive artificial selections than indigenous chickens as generally understood.

Secondly, to reveal the genetic differences between traditional selection and industrial selection, we compared the indigenous chicken group with the commercial chicken group and identified a large number of selective sweeps (6,735) and DSSs (2,532) involving 2,147 genes (Tables S9 and S19). This result suggests that indigenous chickens and the commercial chickens have gone through quite different artificial selection routes as commonly understood.

Thirdly, to reveal uniquely selective sweeps of each chicken breed, we compared each domestic chicken breed with the rest domestic chicken breeds and found that broilers and layers have much higher number of uniquely selective sweeps (7,239 and 6,401, respectively) and DSSs (2,514 and 2,479, respectively) than the indigenous breeds (2,801∼4,555 and 1,363∼1,890, respectively) (Tables S9 and S20∼26).

Fourthly, to reveal possible selective sweeps underlying the miniature body size of Daweishan chicken, we compared Daweishan chicken with the group of Hu chicken, Wuding chicken and Broiler (HWB), with a relatively large body size, and identified 6,359 selective sweeps and 2,263 DSSs including 1,911 genes (Tables S9 and S27).

Finally, to find the selection difference between broilers and layers, we compared the two groups and identified 7,284 selective sweeps and 2,630 DSSs including 1,802 genes (Tables S9 and S28). For the similar comparison in a previous study (Qanbari, et al. 2019), only 41 selective sweeps (40 kb) were identified. Therefore, we identified substantially more selective sweeps. This might be because we use a more rigorous Null model for normalizing *ZF*_*ST*_ and *Δπ*, while the previous study employed a mixture model (Qanbari, et al. 2019).

### Amino-acid altering SNPs are enriched in the selective sweeps

To identify selective sweeps that might be responsible for the formation of a chicken breed, we took the union of DSSs found in comparisons with breed alone as one of the compared group, e.g, for Daweishan chicken, we took the union of DSSs in comparisons Daweishan VS. RJF, Daweishan VS. Other Breeds and Daweishan VS. HWB; and for Hu chicken, we took the union of DSSs in comparisons Hu VS. RJF and Hu VS. Other Breeds; and so on. We identified from 1.22 million (Nine-claw chicken) to 4.06 million (broilers) SNPs in the union of DSSs in each domestic chicken breed (Table 5). Among these SNPs, only 1.28% (Nine-claw chicken) ∼2.02% (Hu chicken) are located in coding regions, while the remaining vast majority (97.98%∼98.72%) fall in non-coding regions (Table 5). As non-coding regions comprise 96.92% of the reference chicken genome (GRCg7b assembly), as in the case of all the bi-allelic SNPs (Table 2), the SNPs in the DSSs are also enriched in non-coding regions relative to in coding regions. Among the SNPs in coding regions, 36.30%∼45.23% are amino-acid altering, which are higher than the corresponding values of all the bi-allelic SNPs (33.33%∼38.33%) (Table 2), suggesting that amino-acid altering SNPs are enriched in the selective sweeps relative to all the bi-allelic SNPs (p = 0.005, K-S test).

**Table 5:** Functional annotation of SNPs in DSSs of each domestic chicken breed

|  | Daweishan | Hu | Piao | Wuding | Nine-claw | Broiler | Layer |
| --- | --- | --- | --- | --- | --- | --- | --- |
| # Total bi-allelic SNPs | 3,657,983 | 1,919,925 | 2,997,593 | 3,020,337 | 1,219,065 | 4,058,599 | 2,862,137 |
| # SNPs in intergenic region (%) | 1,237,910 (33.84) | 662,275 (34.49) | 1,026,302 (34.24) | 982,969 (32.55) | 424,878 (34.85) | 1,279,806 (31.53) | 890,623 (31.12) |
| # SNPs in Intronic region (%) | 1,753,832 (47.95) | 914,031 (47.61) | 1,420,985 (47.40) | 1,491,637 (49.39) | 587,664 (48.21) | 2,030,548 (50.03) | 1,455,882 (50.87) |
| # SNPs in up/downstream (%) | 115,633 (3.16) | 66,121 (3.44) | 98,454 (3.28) | 99,913 (3.31) | 32,073 (2.63) | 108,623 (2.68) | 75,384 (2.63) |
| # SNPs in UTR 3'/UTR 5' (%) | 137,075 (3.75) | 68,060 (3.54) | 106,584 (3.56) | 112,934 (3.74) | 40,400 (3.31) | 151,470 (3.73) | 101,979 (3.56) |
| # SNPs in splicing region (%) | 323 (0.01) | 163 (0.01) | 248 (0.01) | 270 (0.01) | 86 (0.01) | 312 (0.01) | 196 (0.01) |
| # SNPs in ncRNA (%) | 347,114 (9.49) | 170,583 (8.88) | 285,264 (9.52) | 276,744 (9.16) | 118,388 (9.71) | 431,263 (10.63) | 299,387 (10.46) |
| # SNPs in coding region (%) | 66,096 (1.81) | 38,692 (2.02) | 59,756 (1.99) | 55,870 (1.85) | 15,576 (1.28) | 56,577 (1.39) | 38,686 (1.35) |
| # Nonsynonymous-INTOL SNPs (%) | 2,966 (0.08) | 1,620 (0.08) | 2,898 (0.10) | 2,445 (0.08) | 586 (0.05) | 2,247 (0.06) | 1,530 (0.05) |
| # Nonsynonymous-TOL SNPs (%) | 25,045 (0.68) | 15,125 (0.79) | 23,725 (0.79) | 21,523 (0.71) | 5,561 (0.46) | 18,136 (0.45) | 12,407 (0.43) |
| # Synonymous SNPs (%) | 37,058 (1.01) | 21,352 (1.11) | 32,281 (1.08) | 31,094 (1.03) | 9,332 (0.77) | 35,823 (0.88) | 24,429 (0.85) |
| # Stop-gain/loss (%) | 427 (0.01) | 252 (0.01) | 409 (0.01) | 371 (0.01) | 73 (0.01) | 252 (0.01) | 204 (0.01) |
| # No prediction (%) | 600 (0.02) | 343 (0.02) | 443 (0.01) | 437 (0.01) | 24 (0.00) | 119 (0.00) | 116 (0.00) |

### Our predicted selective sweeps are supported by experimental data

To evaluate our detected selective sweeps, we first compared them (Tables S10∼S28) with the 15,439 QTLs in the chicken QTL database (Hu, et al. 2022). We find that 90.5%∼98.3% putative DSSs in each of our comparisons overlap one or more QTLs in the chicken QTLdb (Tables 6 and S29), suggesting that our approach of finding selective sweeps achieves high precision (or positive prediction values). On the other hand, we find that 23.9%∼41.6% QTLs in chicken QTLdb overlap one or more our predicted DSSs in each of our comparisons (Tables 6 and S29), and 11,449 (74.2%) QTLs in the chicken QTLdb overlap one or more of our predicted DSSs in different comparisons, suggesting that our approach of finding selective sweeps is quite sensitive.

**Table 6:** Summary of putative DSSs overlapped with chicken QTLs

| Breeds | # Putative DSSs | # Putative DSSs overlap QTLs in chicken QTLdb (%) | # QTLs in chicken QTLdb overlap our putative DSSs (%) |
| --- | --- | --- | --- |
| Daweishan VS. RJF | 1,550 | 1,472 (95.0%) | 5,526 (35.8%) |
| Hu VS. RJF | 1,811 | 1,702 (94.0%) | 5,247 (34.0%) |
| Piao VS. RJF | 1,483 | 1,342 (90.5%) | 4,801 (31.1%) |
| Wuding VS. RJF | 2,077 | 1,979 (95.3%) | 5,381 (34.9%) |
| Nine-claw VS. RJF | 1,073 | 990 (92.3%) | 3,686 (23.9%) |
| Broiler VS. RJF | 2,745 | 2,601 (94.8%) | 5,463 (35.4%) |
| Layer VS. RJF | 1,926 | 1,837 (95.4%) | 5,334 (34.5%) |
| Indigenous VS. RJF | 1,319 | 1,266 (96.0%) | 5,010 (32.5%) |
| Commercial VS. RJF | 2,382 | 2,237 (93.9%) | 5,086 (32.9%) |

As an additional validation of our detected selective sweeps, we next compared the genes in our predicted DSSs with those that have been reported to be under selection during chicken domestication process, and we describe a few examples here. It has been shown that the *BCO2* locus is involved in the yellow skin trait in domestic chickens (Eriksson, et al. 2008), and we confirmed this results in several of our comparisons, including Daweishan VS. RJF (*F*_*ST*_ support, *ZF*_*ST*_ = 11.27), Daweishan VS. HWB (*F*_*ST*_ support, *ZF*_*ST*_ = 14.79), Hu VS. RJF (Two methods support, *ZF*_*ST*_ = 18.26, *Z*|*Δπ*| = 4.86), Hu VS. Other Breeds (*F*_*ST*_ support, *ZF*_*ST*_ = 6.27), Piao VS. Other Breeds (*F*_*ST*_ support, *ZF*_*ST*_ = 29.36), Wuding VS. Other Breeds (*F*_*ST*_ support, *ZF*_*ST*_ = 22.18), Broiler VS. RJF (*Δπ* support, *Z*|*Δπ*| = 5.08), Broiler VS. Layer (*F*_*ST*_ support, *ZF*_*ST*_ = 9.90), Layer VS. RJF (Two methods support, *ZF*_*ST*_ = 15.66, *Z*|*Δπ*| = 3.17), Layer VS. Other Breeds (*F*_*ST*_ support, *ZF*_*ST*_ = 8.70), Indigenous VS. Commercial (Two methods support, *ZF*_*ST*_ = 26.75, *Z*|*Δπ*| = 3.67) and Commercial VS. RJF (Two methods support, *ZF*_*ST*_ = 27.16, *Z*|*Δπ*| = 4.44) (Figures 2, 3, S2 and S3). The *TSHR* locus is known to be involved in regulation of reproduction and metabolic functions in commercial chickens (Rubin, et al. 2010), and we found that the locus was in selective sweeps in comparisons Hu VS. Other Breeds (*F*_*ST*_ support, *ZF*_*ST*_ = 11.75) and Indigenous VS. RJF (*F*_*ST*_ support, *ZF*_*ST*_ = 13.43) (Figures 2 and S2). It has been reported that the *HNF4G* and *IGF1* loci are involved in growth regulation in chicken (Rubin, et al. 2010; Elferink, et al. 2012), we found that the two loci were in selective sweeps in comparisons Broiler VS. RJF (For *HNF4G, F*_*ST*_ support, *ZF*_*ST*_ = 24.27; For *IGF1, F*_*ST*_ support, *ZF*_*ST*_ = 32.54), Commercial VS. RJF (For *HNF4G, F*_*ST*_ support, *ZF*_*ST*_ = 26.25; For *IGF1, F*_*ST*_ support, *ZF*_*ST*_ = 15.51) and Indigenous VS. Commercial (For *HNF4G, F*_*ST*_ support, *ZF*_*ST*_ = 24.51; For *IGF1, F*_*ST*_ support, *ZF*_*ST*_ = 17.86) (Figures 2 and S2). It has been reported that the *PMCH* locus is related to regulation of appetite and metabolic functions (Shimada, et al. 1998; Rubin, et al. 2010), and we found that the locus was in the selective sweeps in several of our comparisons including Daweishan VS. RJF (*Δπ* support, *Z*|*Δπ*| = 3.25), Daweishan VS. HWB (*Δπ* support, *Z*|*Δπ*| = 6.04), Wuding VS. RJF (*Δπ* support, *Z*|*Δπ*| = 4.82), Wuding VS. Other Breeds (*Δπ* support, *Z*|*Δπ*| = 4.07), Broiler VS. RJF (Two methods support, *ZF*_*ST*_ = 32.54, *Z*|*Δπ*| = 6.05), Broiler VS. Other Breeds (Two methods support, *ZF*_*ST*_ = 19.59, *Z*|*Δπ*| = 6.34), Broiler VS. Layer (*F*_*ST*_ support, *ZF*_*ST*_ = 27.21), Commercial VS. RJF (*F*_*ST*_ support, *ZF*_*ST*_ = 15.51) and Indigenous VS. Commercial (*F*_*ST*_ support, *ZF*_*ST*_ = 17.86) (Figures 2, 3, S2 and S3). It has been shown that the *INSR* locus is related to the growth of chicken by encoding a critical component in insulin signaling (Rubin, et al. 2010), and we found that the locus was in the selective sweeps in comparisons Daweishan VS. HWB (Two methods support, *ZF*_*ST*_ = 7.37, *Z*|*Δπ*| = 4.46), Broiler VS. RJF (Two methods support, *ZF*_*ST*_ = 15.30, *Z*|*Δπ*| = 5.23), Broiler VS. Other Breeds (Two methods support, *ZF*_*ST*_ = 11.61, *Z*|*Δπ*| = 4.86) and Broiler VS. Layer (*F*_*ST*_ support, *ZF*_*ST*_ = 9.78) (Figures 2, 3, S2 and S3). It has been shown that the *NELL1* locus is related to the skeletal integrity in broiler (Elferink, et al. 2012), and we found that the locus was in the selective sweeps of the Broiler VS. Other Breeds comparison (*F*_*ST*_ support, *ZF*_*ST*_ = 22.06) (Figure S2). It has been reported that the *IRX4* locus is related to the rumpless trait of Piao chicken (Wang, Khederzadeh, et al. 2021), and we found that the locus was in the selective sweeps in comparisons Piao VS. RJF (*F*_*ST*_ support, *ZF*_*ST*_ = 13.79) and Piao VS. Other Breeds (*F*_*ST*_ support, *ZF*_*ST*_ = 12.12) (Figures 2 and S2). The other selective sweep loci found in the previous studies are also confirmed by our results, such as *ALX1, KITLG, EGFR, DLK1, JPT2* (annotated as *HN1L* in GRCg7b), *CRAMP1* and *GLI3* loci, which are related to the general domestication process of chicken (Qanbari, et al. 2019), the *SKIV2L2* locus that is related to pigmentation (Qanbari, et al. 2019), and the *LEPR, MEGF10* and *SPEF2* loci, which are related to production-oriented selection (Qanbari, et al. 2019) (Figures 2, 3, S2 and S3). Taken together, all these results suggest that our approach is highly reliable to find selective sweeps in domestic chickens.

### Novel selective sweeps are found in the chicken breeds

In addition to confirming many previously identified selective sweeps containing genes related to chicken domestication as described above, we also find numerous novel selective sweeps containing genes (Tables S11∼S28) or in gene deserts. We now highlight a few of them with extremely high *ZF*_*ST*_ and/or *Z*|*Δπ*| values in each comparison (Figures 2, 3, S2 and S3). Gene *ARHGAP39* on chromosome 2 is in the selective sweeps with extremely high *ZF*_*ST*_ and/or *Z*|*Δπ*| value in comparisons Daweishan VS. RJF, Piao VS. RJF, Nine-claw VS. RJF, Indigenous VS. RJF, Nine-claw VS. Other Breeds and Broiler VS. RJF. *ARHGAP39* plays important roles in cell cytoskeletal organization, growth, differentiation, neuronal development and synaptic functions (Moon and Zheng 2003). Gene *TIGD5* on chromosome 2 is in the selective sweeps with extremely high *Z*|*Δπ*| value in comparisons Daweishan VS. RJF, Piao VS. RJF, Daweishan VS. Other Breeds, Hu VS. Other Breeds, Piao VS. Other Breeds, Wuding VS. Other Breeds, Daweishan VS. HWB and Indigenous VS. Commercial. *TIGD5* encodes the tigger transposable element-derived protein 5 and is important for nucleic acid binding (Nusbaum, et al. 2006). Gene *KCNK16* on chromosome 3 is in the selective sweeps with extremely high *ZF*_*ST*_ value in comparisons Layer VS. Other Breeds and Broiler VS. Layer. *KCNK16* encodes a rapidly activating and non-inactivating outward rectifier K^+^ channel (Girard, et al. 2001). Gene *CD8A* on chromosome 4 is in the selective sweeps with extremely high *Z*|*Δπ*| value in comparisons Wuding VS. RJF, Layer VS. RJF, Commercial VS. RJF, Daweishan VS. Other Breeds, Piao VS. Other Breeds, Wuding VS. Other Breeds, Nine-claw VS. Other Breeds, Broiler VS. Other Breeds, Layer VS. Other Breeds, Indigenous VS. Commercial and Daweishan VS. HWB. *CD8A* encodes the T-cell surface glycoprotein CD8 alpha chain precursor and plays essential roles in immune response (Littman, et al. 1985). Gene *COL6A2* on chromosome 7 is in the selective sweeps with extremely high *ZF*_*ST*_ value in comparisons Daweishan VS. Other Breeds, Daweishan VS. HWB and Piao VS. Other Breeds. The gene encodes the collagen alpha-2(VI) chain precursor which act as a cell-binding protein (Gerhard, et al. 2004). Besides the genes mentioned above, we also indicate in Figures 2, 3, S2 and S3 many other genes located in novel selective sweeps with extremely high *ZF*_*ST*_ and/or *Z*|*Δπ*| values in multiple comparisons such as: *ANXA10* on chromosome 3, gene *LOC*107053954 on chromosome 8, gene *SLC16A9* on chromosome 6, gene *ENS-1* on chromosome W and gene *HSD17B4* on chromosome Z. It is interesting to experimentally investigate the roles of these genes in the domestication and breeding of each chicken breed.

In Figures 2, 3, S2 and S3, we also label a few examples of selective sweeps in gene desserts, with extremely high *ZF*_*ST*_ and/or *Z*|*Δπ*| values in multiple comparisons. It is highly likely that these selective sweeps might harbor non-coding functional sequences such as *cis*-regulatory modules of distal genes.

### Selective sweeps related to each chicken breed

In addition to finding numerous novel selective sweeps containing genes in each comparison (Tables S11∼S28), we also identify numerous unique selective sweeps/DSSs that are only seen in comparisons with a breed alone as one of the two compared groups or selective sweeps/DSSs containing genes with interesting functions. These selective sweeps/DSSs might contain genes (Tables S30∼S36) whose functions are related to the specific traits of the chicken breed. Specifically, for Daweishan chicken, we identified 44 putative genes in the selective sweeps that might be related to its unique traits including the small body size (Table S30). For example, the *GHR* (growth hormone receptor) gene is located in a selective sweep window on chromosome Z, which overlaps body weight QTLs and shank length QTLs. The gene is in the selective sweep windows identified in comparisons with Daweishan chicken alone as one of the two compared groups (Daweishan VS. RJF, *ZF*_*ST*_ = 17.71; Daweishan VS. Other Breeds, *ZF*_*ST*_ = 17.69; Daweishan VS. HWB, *ZF*_*ST*_ = 19.51) (Figures 2 and S2). It has been reported that loss-of-function mutations in *GHR* was related to sex-linked dwarfism in chicken (Agarwal, et al. 1994). We analyzed the SNPs in the *GHR* gene body for each chicken breeds using the GRCg7b assembly as the template and found 79 unique SNPs in the gene of Daweishan chicken, which were not present in the other chicken breeds (Hu, Piao, Wuding, Nine-claw, Broiler, Layer and RJF). Among these 79 unique SNPs, 68 are in intronic regions, 10 are in UTRs and one is nonsynonymous SNP that leads to a CGG to TGG (R to W) mutation, which is tolerant. The substitution allele has a frequency of 0.76, thus it is only nearly fixed. As no fixed potential amino acid-altering mutation could be found in the *GHR* coding regions, we hypothesize that *GHR* gene might be related to the small body size of Daweishan chicken through changes in its regulatory sequences in the window, resulting in a decrease in its expression. This possibility warrants further experimental investigation.

For Hu chicken, we identified 14 putative genes in selective sweeps (Table S31) that might be related to its unique traits including the very stout legs. Specifically, gene *TRIM13* in a selective sweep on chromosome 1 (Hu VS. RJF, *ZF*_*ST*_ = 7.49 and *Z*|*Δπ*| = 7.19; Hu VS. Other Breeds, *ZF*_*ST*_ = 10.01 and *Z*|*Δπ*| = 4.58) (Figures 2, 3, S2 and S3) overlaps shank circumference QTLs, and there are two nonsynonymous SNPs in the gene body which are fixed (allele frequency = 1). Thus, it is interesting to experimentally investigate the role of *TRIM13* in the development of the very stout legs of Hu chicken.

For Piao chicken, we identified six putative genes in selective sweeps (Table S32) that might be related its unique traits including the rumpless trait. Of these six genes, *IRX4* in a selective sweep on chromosome 2 was reported to be related to the rumpless trait of Piao chicken in a previous study (Wang, Khederzadeh, et al. 2021), and we also found that the selective sweeps were only identified by the comparisons Piao VS. RJF (*ZF*_*ST*_ = 13.79) and Piao VS. Other Breeds (*ZF*_*ST*_ = 12.12) (Figures 2 and S2, Tables S12 and S23). Thus, it is highly likely that *IRX4* is related to the rumpless trait of Piao chicken. At the same time, the previous study also identified genes *IL18, HSPB2*, and *CRYAB* to be related to the rumpless trait of Piao chicken. Although we also found these three genes in the selective sweeps for the comparison Piao VS.

Other Breeds (Table S23 and Figure S2), these three genes were also present in the selective sweeps for comparisons with a breed having normal tails alone as one of the two compared groups, such as Daweishan VS. RJF (Table S10, Figures 2 and S2), Hu VS. RJF (Table S11, Figures 2 and S2), Nine-claw VS. RJF (Table S14, Figures 2 and S2), Broiler VS. RJF (Table S15, Figures 2 and S2), Layer VS. RJF (Table S16, Figure 2) and Daweishan VS. HWB (Table S21, Figure 2). Therefore, these three genes might not be related to the rumpless trait of Piao chicken.

For Wuding chicken, we identified 18 putative genes in selective sweeps (Table S33) that might be related to its unique traits including colorful feathers and thick fat. Specifically, gene *SSTR5* in a selective sweep on chromosome 14 (Wuding VS. RJF, *ZF*_*ST*_ = 7.38 and *Z*|*Δπ*| = 5.16; Wuding VS. Other Breeds, *Z*|*Δπ*|| = 3.33) (Figures 2, 3 and S3) overlaps body weight QTLs, however, there are no nonsynonymous SNPs in its gene body. Gene *LOC*101748311 in a selective sweep on chromosome 1 (Wuding VS. Other Breeds comparison, *ZF*_*ST*_ = 9.70 and *Z*|*Δπ*| = 3.59) (Figures S2 and S3) overlaps the feather density QTLs and comb length QTLs and there are two nonsynonymous SNPs in its gene body, but their allele frequencies are very low (< 0.2). It is likely that both genes might be related to Wuding chicken’s traits by changes in regulatory regions, which warrants further experimental studies.

For Nine-claw chicken, we identified seven putative genes in selective sweeps (Table S34) that might be related to its unique traits. Specifically, gene *ATG4B* on chromosome 9 (Nine-claw VS. RJF, *ZF*_*ST*_ = 7.00 and *Z*|*Δπ*| = 3.60; Nine-claw VS. Other Breeds, *ZF*_*ST*_ = 7.56) (Figures 2, 3 and S2) overlaps egg production rate QTLs, but there are no nonsynonymous SNPs in its gene body.

For Broilers, we have identified 151 putative genes in selective sweeps (Table S35) that might be related to its unique traits including the fast growth rate. Of these genes, *GHRHR* on chromosome 2 (Growth hormone-releasing hormone receptor) (Broiler VS. Other Breeds, *ZF*_*ST*_ = 10.36; Broiler VS. Layer, *ZF*_*ST*_ = 7.09) (Figure S2) is well-known for its role in determining growth rate and body size via regulating the growth hormone (GH) level in blood (Jia, et al. 2018), however, there are no nonsynonymous SNPs in its gene body; *IGF2BP1* on chromosome 27 (insulin-like growth factor 2 mRNA-binding protein 1) (Broiler VS. RJF, *ZF*_*ST*_ = 7.69 and *Z*|*Δπ*| = 3.98) (Figures 2 and 3) may affect growth rate via regulating insulin-like growth factor 2 level (Cao, et al. 2018), but there are no nonsynonymous SNPs in its gene body. The *IGF2BP1* locus also overlaps the claw percentage QTLs, shank length QTLs, claw weight QTLs, drumstick and thigh weight QTLs, breastbone crest length QTLs, body weight QTLs, body slope length QTLs and femur bending strength QTLs. The result is consistent with a recent report that mutations in the promoter region of the *IGF2BP1* gene can affect chicken body size (Wang, Hu, et al. 2021). In addition, the following genes are also interesting as they overlap white striping QTLs, abdominal fat percentage QTLs, wooden breast QTLs and body weight QTLs, and thus might be related to the large body and fast growth rate of broilers, including *OVALX* on chromosome 2 (Broiler VS. RJF, *ZF*_*ST*_ = 9.41 and *Z*|*Δπ*| = 5.78; Broiler VS. Other Breeds, *ZF*_*ST*_ = 8.45 and *Z*|*Δπ*| = 4.18) (Figures 2, 3, S2 and S3), *RRAS* on chromosome 5 (Broiler VS. RJF, *ZF*_*ST*_ = 14.87 and *Z*|*Δπ*| = 7.53; Broiler VS. Other Breeds, *ZF*_*ST*_ = 11.73 and *Z*|*Δπ*| = 6.00) (Figures 2, 3, S2 and S3), *SYT8* on chromosome 5 (Broiler VS. RJF, *ZF*_*ST*_ = 6.90 and *Z*|*Δπ*| = 5.87; Broiler VS. Other Breeds, *ZF*_*ST*_ = 6.89 and *Z*|*Δπ*| = 3.85) (Figures 2, 3, S2 and S3), *TNNI2* on chromosome 5 (Broiler VS. RJF, *ZF*_*ST*_ = 6.90 and *Z*|*Δπ*| = 5.87; Broiler VS. Other Breeds, *ZF*_*ST*_ = 6.89 and *Z*|*Δπ*| = 3.85) (Figures 2, 3, S2 and S3) and *FBXO28* on chromosome 3 (Broiler VS. RJF, *ZF*_*ST*_ = 10.25 and *Z*|*Δπ*| = 5.30; Broiler VS. Other Breeds, *ZF*_*ST*_ = 8.38 and *Z*|*Δπ*| = 4.90) (Figures 2, 3, S2 and S3). All these genes either has no nonsynonymous SNPs or the allele frequencies of the nonsynonymous SNPs are very low. Thus, it is highly likely that they might be related to the broilers’ traits through changes in their regulatory regions.

For layers, we identified 36 genes in selective windows (Table S36) that might be related to its unique traits including larger number of egg-production. Specifically, gene *NOCT* (Nocturnin), *ELF2* (ETS-related transcription factor Elf-2) and *MGARP* (mitochondria localized glutamic acid rich protein) are all located in the same selective sweep on chromosome 4 (Layer VS. RJF, *ZF*_*ST*_ = 9.84 and *Z*|*Δπ*| = 4.61; Layer VS. Other Breeds, *ZF*_*ST*_ = 10.80 and *Z*|*Δπ*| = 5.24; Broiler VS. Layer, *ZF*_*ST*_ = 9.23 and *Z*|*Δπ*| = 5.20) (Figures 2, 3, S2 and S3). *NOCT* is expressed in retina and many other tissues, and its expression shows circadian rhythm (Baggs and Green 2003; Stubblefield, et al. 2012). *NOCT* is known to be involved in adipogenesis, osteogenesis, and obesity in mice (Hughes, et al. 2018). It has been shown that *MGARP* is involved in the synthesis of estrogen in ovary, and its expression is under the control of the hypothalamic-pituitary-gonadal (HPG) axis (Zhou, et al. 2011). It has been reported that *ELF2* plays a role in cell proliferation (Chung, et al. 2016). Moreover, gene *SLC25A15* on chromosome 1 (mitochondrial ornithine transporter 1) (Layer VS. RJF, *ZF*_*ST*_ = 7.23 and *Z*|*Δπ*| = 3.96; Layer VS. Other Breeds, *ZF*_*ST*_ = 9.68 and *Z*|*Δπ*| = 3.82; Broiler VS. Layer, *ZF*_*ST*_ = 10.52 and *Z*|*Δπ*| = 3.21) (Figures 2, 3, S2 and S3) overlaps the oviduct length QTLs, thus might be related to the egg-production rate of layers. Thus, these four genes might be related to the layers’ unique traits. However, the four genes either have no nonsynonymous SNPs or the allele frequencies of nonsynonymous SNPs are very low. It is interesting to experimentally investigate roles of variations in the regulatory regions of these genes in the high egg-production related traits of layers, such as the lack of brooding behaviors, egg-laying circadian rhythm and high demand for light.

## Discussion

Next generation sequencing technology makes it possible to re-sequence a large number of individuals for a species for genome-wide studies. In 2021, NCBI released more complete domestic chicken (*Gallus gallus*) genome assemblies (GRCg7b and GRCg7w), providing good reference genomes for this economically, medically and evolutionally important bird. Using the GRCg7b assembly as the template, we have called the variants in populations of eight chicken breeds including 25 Daweishan chickens, 10 Hu chickens, 23 Piao chickens, 23 Wuding chickens, four Nine-claw chickens, 60 broilers, 56 layers and 35 RJFs. By comparing the putative selective sweeps of Daweishan, Hu, Piao, Wuding, Nine-claw chicken, broilers and layers with respect to others and RJFs (19 comparisons, Tables 4 and S9), we identified putative selective sweeps and genes that might be related to the specific-traits of each chicken breed or groups (Tables S30∼36). Remarkably, the vast majority (90.5%∼98.3%) of our identified DSSs in each of our 19 comparisons overlap QTLs in the chicken QTLdb (Tables 6 and S29), while 74.2% QTLs in the chicken QTLdb overlap our identified DSSs in different comparisons. Moreover, we also confirm many previously identified genes under artificial selection. Thus, we have achieved very high prediction precision (or positive prediction values) and sensitivity.

More importantly, our analyses also result in many new findings. Firstly, we identify a much larger number of selective sweeps/DSSs and genes related to the specific traits of broilers and layers than the previous study (Qanbari, et al. 2019). We attribute the difference to the different statistic models used in the two studies. More specifically, we use a more rigorous Null model by generating 100-sets of windows with the allele frequencies randomly permutated (Churchill and Doerge 1994). Using the mean and standard deviation of the Null model, we compute *ZF*_*ST*_ and *Z*|*Δπ*| for each window in each comparison. In contrast, the previous study (Qanbari, et al. 2019) used the mean and standard deviation of the *F*_*ST*_ and |*Δπ*| values of the windows to compute the *ZF*_*ST*_ and *Z*|*Δπ*|, which is not a rigorous Null model. Thus, the previous study might underestimate the number of selective sweeps. Consequently, we identify ∼2,500 putative DSSs containing ∼1,800 genes for the broilers and ∼2,000 putative DSSs containing ∼1,000 genes for the layers (Tables 4 and S9), which included almost all the only 90 and 66 putative selective sweep windows (40 kb) found in broilers and layers, respectively, in the previous study (Qanbari, et al. 2019).

Secondly, we negate several genes found in a previous study (Wang, Khederzadeh, et al. 2021) to be related to the rumpless trait of Piao chicken based on our results from multiple comparisons with or without Piao chicken. More specifically, in addition to *IRX4*, the previous study also claimed that *IL18, HSPB2*, and *CRYAB* (Wang, Khederzadeh, et al. 2021) might be related to the rumpless trait of Piao chicken. Although we also find that gene *IRX4* presents in putative selective sweeps only in the Piao VS. RJF and Piao VS. Other Breeds comparisons (Tables S12 and S23), thus it might be related to the rumpless trait of Piao chicken. However, genes *IL18, HSPB2*, and *CRYAB* present in selective sweeps in not only the comparison related to Piao chicken (Piao VS. Other Breeds, Table S23), but also in comparisons with chickens having a normal tail alone as a group, such as Daweishan, Wuding, Nine-claw chicken, broilers and layers (Tables S10, S11, S14, S15, S16 and S21). Thus, these three genes might not be related to the rumpless trait of Piao chicken.

Thirdly, our analyses provide many novel selective sweeps containing genes that might be related to artificial selection of unique traits of each chicken breed (Tables S30∼S36), and some are quite interesting, thus warranting further experimental investigations. For example, it is interesting to test the roles of the genes *NOCT* and *MGARP* in the high egg-production of layers.

Finally, we find that although SNPs in selective sweeps are more likely to alter amino acids than expected, many genes in selective sweeps often lack fixed amino acid-altering mutations. These genes might affect the traits of chicken breeds by changing their expression levels through changes in their cis-regulatory regions. Consistently with this argument, we found that SNPs in non-coding regions in general, or in selective sweeps in particular, are enriched in all the eight chicken breeds analyzed in this study. Due to the lack of a more complete map of *cis*-regulatory modules and their constituent transcription factor binding sites in the chicken reference genome, it is difficult to further pin down the sites that affect the expression of these genes and related organism traits. Therefore, it is pressing to map out the *cis*-regulatory elements in the chicken reference genome as has been done for *C. elegans*(Van Nostrand and Kim 2013; Kudron, et al. 2018), *D. melanogaster* (Negre, et al. 2011; Kudron, et al. 2018), mice (Shen, et al. 2012; Moore, et al. 2020) and humans(Gerstein, et al. 2012; Moore, et al. 2020).

## Materials and methods

### Re-sequencing short reads from NCBI SRA

We downloaded genomic short reads for two broiler lines from NCBI Sequence Read Archive (SRA): “Broiler A” (n=40, access number PRJEB15276) and “Broiler B” (n=20, access number PRJEB30270). Broiler A was originally from France, and Broiler B was from the company Indian River International. We downloaded DNA short reads for three layer lines from NCBI SRA: Layer A” (n=25, access number PRJEB15189) were white egg layers, “Layer B” (n=25, access number PRJEB30270) were brown egg layers, and “Layer C” (n=6, access number PRJEB30270) were crossbred layers. We downloaded genomic short reads for two RJF populations from NCBI SRA: “RJF A” (n=25, access number PRJEB3027) were from northern Thailand, and “RJF B” (n=10, access number PRJEB3027) were from India.

### Re-sequencing of indigenous chicken samples

We re-sequenced 85 indigenous chicken individuals from the Experimental Breeding Chicken Farm of the Yunnan Agricultural University (Yunnan, China), including 25 Daweishan chickens aged 10 months (nine males, 16 females), 10 Hu chickens aged seven months (five males, five females), 23 Piao chickens aged 10 months (11 males, 12 females), 23 Wuding chickens aged 10 months (11 males, 12 females) and four Nine-claw chickens aged 10 months (two males, two females).

### Short-reads DNA sequencing

Two milliliters of blood were drawn from the wing vein of each chicken in a centrifuge tube containing anticoagulant (EDTA-2K) and stored at -80°C until use. Genomic DNA (10µg) in each blood sample was extracted using a DNA extraction kit (DP326, TIANGEN Biotech, Beijing, China) and fragmented using a Bioruptor Pico System (Diagenode, Belgium). DNA fragments around 350 bp were selected using SPRI beads (Beckman Coulter, IN, USA). DNA-sequencing libraries were prepared using Illumina TruSeq® DNA Library Prep Kits (Illumina, CA, USA) following the vendor’s instructions. The libraries were subject to 150 cycles paired-end sequencing on an Illumina Novaseq 6000 platform (Illumina, CA, USA) at ∼30 coverage. Variant calling We mapped the short reads of each individual chicken to the reference genome (GRCg7b) using BWA (0.7.17) (Li and Durbin 2009) and SAMtools (1.9) (Li, et al. 2009) with the default settings and called variants for each individual using GATK-HaplotypeCaller (4.1.6) (McKenna, et al. 2010) with the default settings. After generating the GVCF files for each individual, we computed allele frequencies in the same chicken breed using the GATK-CombineGVCFs (4.1.6) tool (McKenna, et al. 2010). For each chicken breed, we removed variants with Quality by depth (QD) < 2, Fisher strand (FS) > 60, Root mean square mapping quality (MQ) < 40, Strand odd ratio (SOR) > 3, Rank Sum Test for mapping qualities (MQRankSum) < -12.5 and Rank Sum Test for site position within reads (ReadPosRankSum) < -8 for SNPs and QD < 2, FS > 200, SOR > 10, Likelihood-based test for the consanguinity among samples (InbreedingCoeff) < -0.8 and ReadPosRankSum < -20 for indels.

### Functional annotation of variants

We used the package ANNOVAR (Wang, et al. 2010) to annotate the variants according to their locations on the reference genome into seven categories including 1) intergenic regions, 2) intronic regions, 3) coding regions (synonymous, nonsynonymous, stop gain and stop loss), 4) up/downstream of a gene, 5) splicing sites, 6) 5’ untranslated regions (5’UTRs) and 3’ UTRs, and 7) non-coding RNA regions. We used the tool Ensembl Variant Effect Predictor (VEP) (McLaren, et al. 2016) to predict the impact of amino acid-altering SNPs.

### Detection of selective sweeps

The selective sweeps were detected using two methods including genetic differentiation (*F*_*ST*_) (Reynolds, et al. 1983) and nucleotide diversity (*π*). We estimated *F*_*ST*_ for each comparison of two chicken populations using VCFtools (0.1.16) (Danecek, et al. 2011) with a sliding window size 40 kb and a step size 20 kb. We estimated *π* for each group using VCFtools (0.1.16) (Danecek, et al. 2011) with a sliding window size 40 kb and a step size 20 kb, and calculated the absolute value of the difference in nucleotide diversity (|*Δπ*|) in each window for each comparison of two populations. For both *F*_*ST*_ and |*Δπ*|, we only used the bi-allelic SNPs on autosomes and sex chromosomes for the analysis. To evaluate the statistic significance of the *F*_*ST*_ and *π* values for a comparison, we generate a Null model by shuffling the allele frequency data for 100 times while keeping the SNP positions fixed (Churchill and Doerge 1994). We then computed *F*_*ST*_ and |*Δπ*| for the permuted windows as well as their means 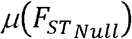 and *μ*(|*Δπ*|_*Null*_)) and standard deviations 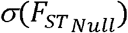 and *σ*(|*Δπ*|_*Null*_)). We computed the *Z* value for each *F*_*ST*_ and |*Δπ*| values for a comparison by using the following formulas:

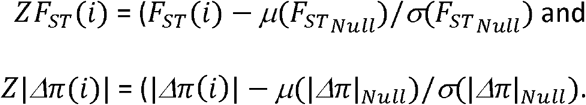

We consider a window with either *ZF*_*ST*_(*i*) > 6 or *Z*|*Δπ*| > 3.09 (P-value < 0.001) to be a putative selective sweep. Since adjacent putative selective sweep windows may overlap with each other, we merged adjacent windows if they overlapped by at least one nucleotide to obtain the discrete selective sweeps (DSSs) for each comparison.

### Selective sweeps analysis

To reveal selective sweeps in the domestic chicken populations, we conducted a total of 19 comparisons, including 1) Daweishan chickens VS. RJF, 2) Hu chickens VS. RJF, 3) Piao chickens VS. RJF, 4) Wuding chickens VS. RJF, 5) Nine-claw chickens VS. RJF, 6) Broilers VS. RJF, 7) Layers VS. RJF, 8) Indigenous chickens (Daweishan chickens + Hu chickens + Piao chickens + Wuding chickens + Nine-claw chickens) VS. RJF, 9) Commercial chickens (layers + broilers) VS. RJF, 10) Indigenous chickens VS. Commercial chickens, 11) Daweishan chickens VS. the other six domestic chicken breeds, 12) Hu chickens VS. the other six domestic chicken breeds, 13) Piao chickens VS. the other six domestic chicken breeds, 14) Wuding chickens VS. the other six domestic chicken breeds, 15) Nine-claw chickens VS. the other six domestic chicken breeds, 16) Broilers VS. the other six domestic chicken breeds, 17) Layers VS. the other six domestic chicken breeds, 18) Daweishan chickens VS. chickens with relatively large body size (Hu chickens, Wuding chickens and Broilers), 19) Broilers VS. Layers.

## Data availability

All the re-sequencing data of the 85 indigenous chickens are available in the SRA database with the accession number of ‘PRJNA893352’ (https://dataview.ncbi.nlm.nih.gov/object/PRJNA893352?reviewer=bkis6iua659tf15ig25ve6g215) and ‘PRJNA865247’ (https://dataview.ncbi.nlm.nih.gov/object/PRJNA865247?reviewer=6q14iagmjc8rtrio5pr2pmf3s6).

## Acknowledgements

This work was supported by the National Natural Science Foundation of China (U2002205 and U1702232), Yunling Scholar Training Program of Yunnan Province (2014NO48), Yunling Industry and Technology Leading Talent Training Program of Yunnan Province (YNWR-CYJS-2015-027), Natural Science Foundation of Yunnan Province (2019IC008 and 2016ZA008), and Department of Bioinformatics and Genomics of the University of North Carolina at Charlotte.

## Notes

### Competing Interest Statement

The authors have declared no competing interest.

