## Supplementary figures for "Artificial selection footprints in domestic chicken genomes"

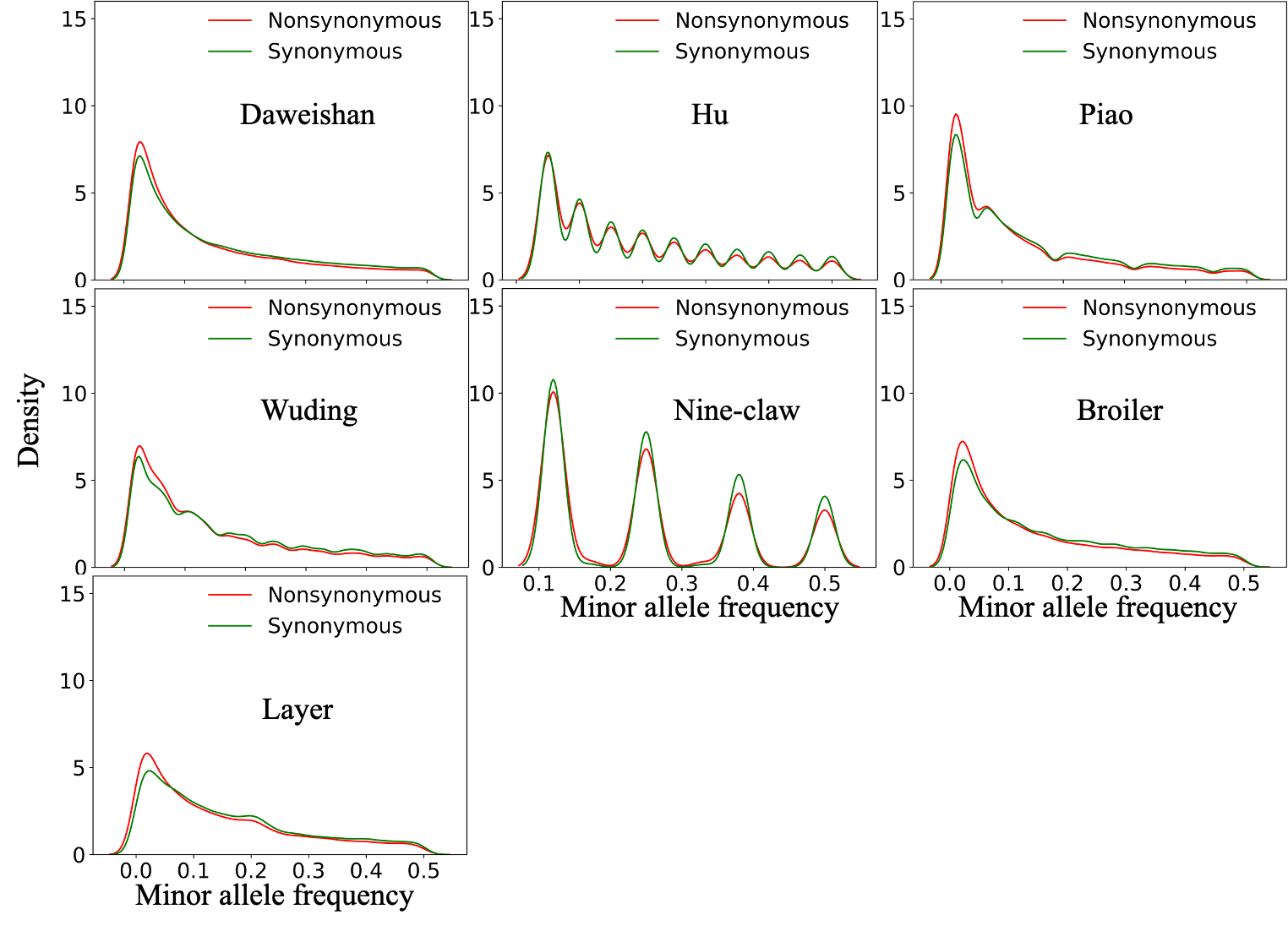

**Figure S1.** Distribution of the minor allele frequency among each chicken breed

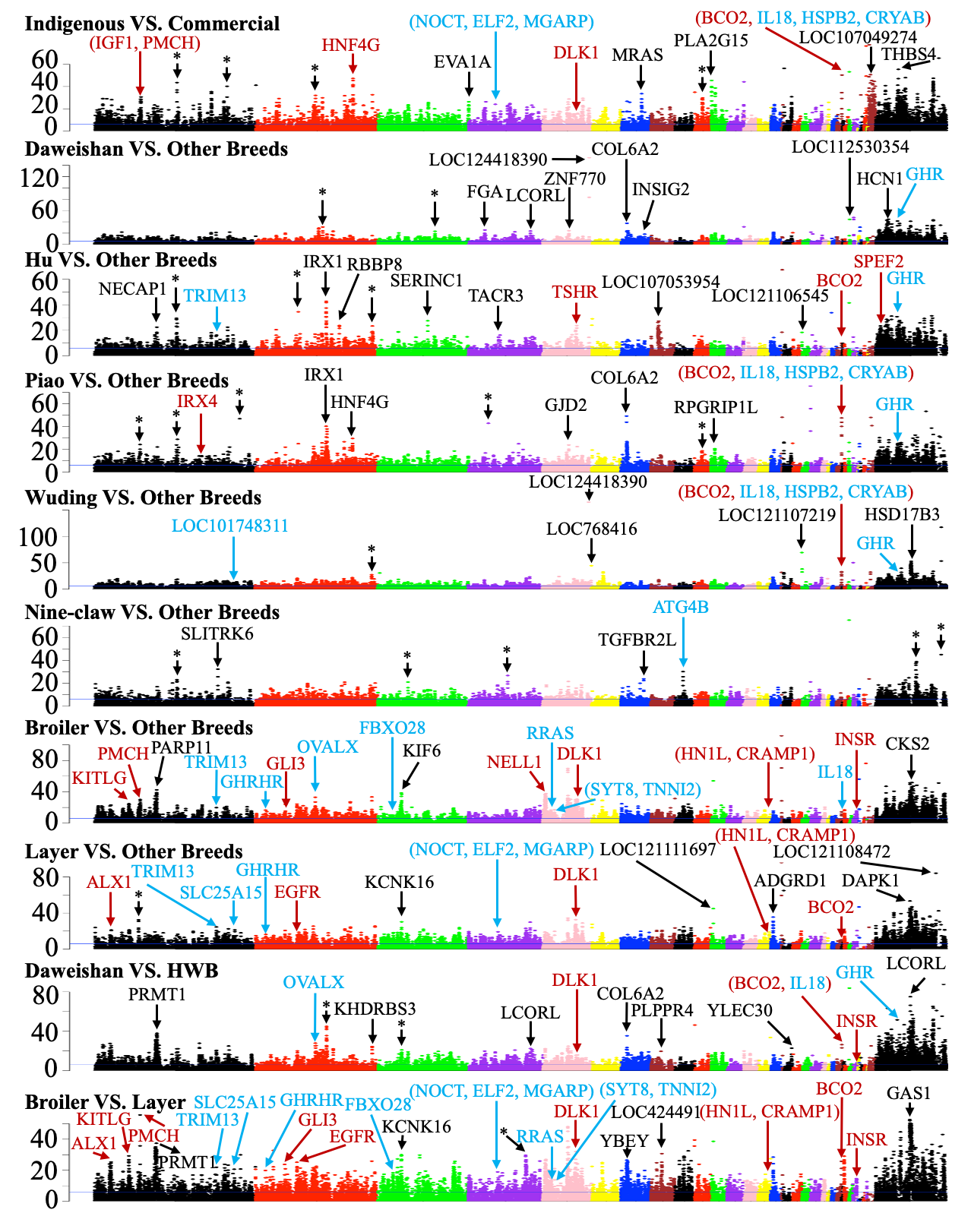

**Figure S2.** Manhattan plots of ${ZF}_{ST}$ values of each window on each chromosome for the indicated comparisons. The blue horizontal line indicates the ${ZF}_{ST}$ cutoff = 6. Examples of genes in significant selective sweep windows are shown in different color. Genes that have been previously reported in selective sweep windows are shown in red, genes in our predicted selective sweep windows potentially related to the specific traits of each chicken breed are shown in blue, and genes in novel selective sweep windows with extremely high ${ZF}_{ST}$ values are shown in black. Asterisk represents selective sweep windows lacking annotated genes.

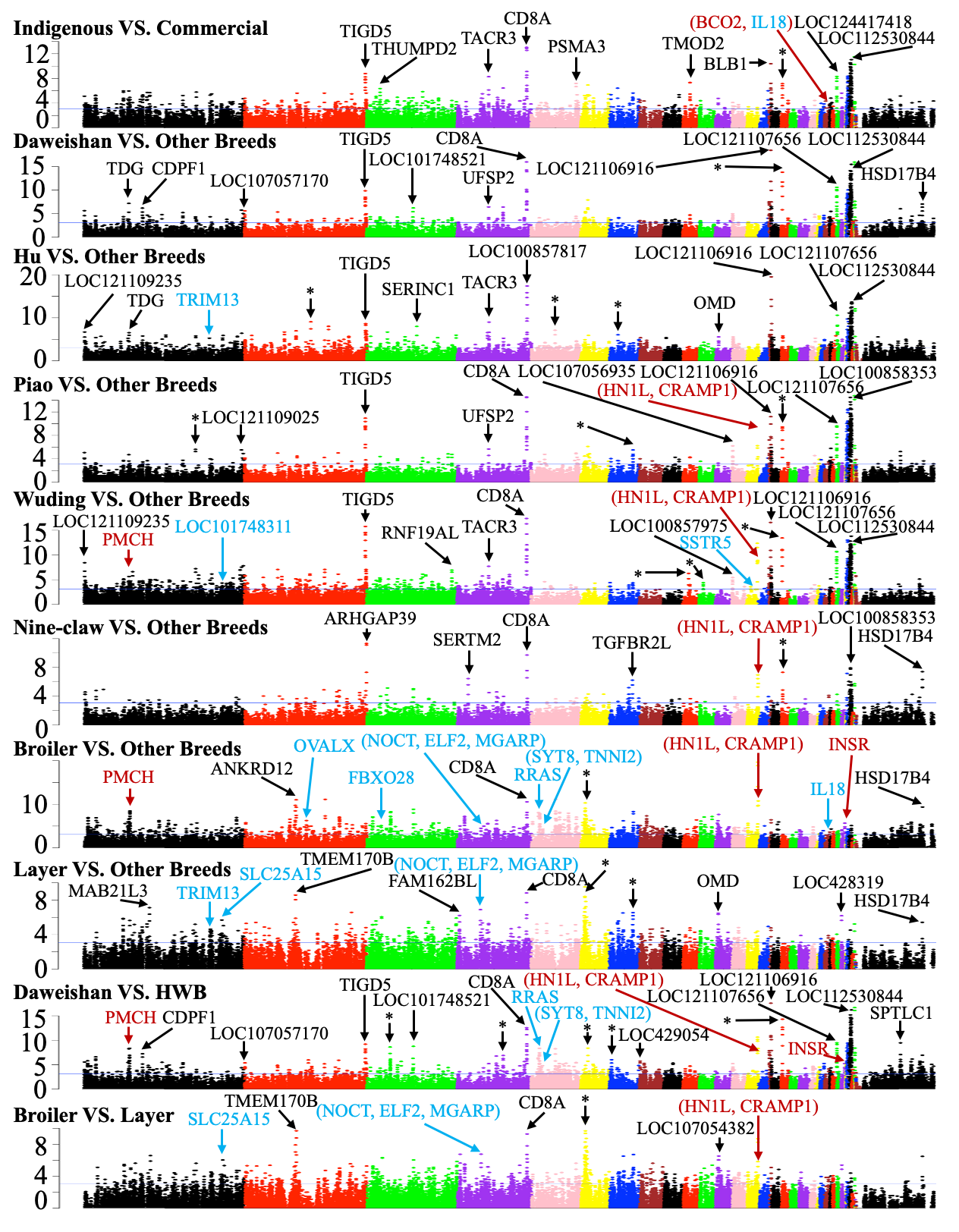

**Figure S3.** Manhattan plots of $Z|\Delta\pi|$ values of each window on each chromosome for the indicated comparisons. The blue horizontal line indicates the $Z|\Delta\pi|$ cutoff = 3.09. Examples of genes in significant selective sweep windows are shown in different color. Genes that have been previously reported in selective sweep windows are shown in red, genes in our predicted selective sweep windows potentially related to the specific traits of each chicken breed are shown in blue, and genes in novel selective sweep windows with extremely high $Z|\Delta\pi|$ values are shown in black. Asterisk represents selective sweep windows lacking annotated genes.
